# Processing of task-irrelevant sounds during typical everyday activities in children

**DOI:** 10.1101/2022.05.19.492605

**Authors:** Ranjan Debnath, Nicole Wetzel

**Affiliations:** Leibniz Institute for Neurobiology, Brenneckestr. 6, 39118 Magdeburg, Germany; Center for Behavioral Brain Sciences, Magdeburg, Germany; University of Applied Sciences Magdeburg-Stendal, Magdeburg, Germany

**Keywords:** Children, Auditory attention, Attention orienting, ERP, Theta power

## Abstract

Our ability to focus on a task and ignore task-irrelevant stimuli is critical for efficient cognitive functioning. Attention control is especially required in the auditory modality as sound has privileged access to perception and consciousness. Despite this important function, little is known about auditory attention during typical everyday activities in childhood. We investigated the impact of task-irrelevant sounds on attention during three everyday activities – playing a game, reading a book, watching a movie. During these activities, environmental novel sounds were presented within a sequence of standard sounds to 7–8-year-old children and adults. We measured ERPs reflecting early sound processing and attentional orienting and theta power evoked by standard and novel sounds during these activities. Playing a game vs. reading or watching reduced early encoding of sounds in children and affected ongoing information processing and attention allocation in both groups. In adults, theta power was reduced during playing at mid-central brain areas. Results show a pattern of immature neuronal mechanisms underlying perception and attention of task-irrelevant sounds in 7–8-year-old children. While the type of activity affected the processing of irrelevant sounds in both groups, early stimulus encoding processes were more sensitive to the type of activities in children.

## 1. Introduction

Children can be easily distracted by task-irrelevant surrounding sounds (Hoyer, Elshafei, Hemmerlin, Bouet, & Bidet-Caulet, 2021; Joseph, Hughes, Sorqvist, & Marsh, 2018; Wetzel, Scharf, & Widmann, 2019). On the other hand, children are sometimes so immersed in an activity that the calls from their parents do not capture their attention. The present study investigates how different everyday life activities affect neuronal processes underlying the perception of task-irrelevant sounds and the allocation of attention toward such sounds in children and an adult control group.

Selective attention, the ability to focus on the task at hand and filter out the task-irrelevant stimuli, is key to efficient cognitive functioning. Although resisting task-irrelevant information is often desirable, novel or unexpected events in the environment even in an unattended sensory modality can involuntarily capture our attention. Such stimulus-driven involuntary shift of attention is part of the orienting response (Sokolov, 1990) and allows evaluation of unexpected events, and is crucial for adaptive behavior. Conversely, the involuntary shift of attention can impair the ongoing cognitive and task-related processes (Escera, Yago, & Alho, 2001; Schröger, Giard, & Wolff, 2000; Schröger & Wolff, 1998). The orienting of attention can be measured using versions of the oddball paradigm that typically include a sequence of repeated standard sounds and rare distractor sounds that deviate in one or more features from standard sounds. Most oddball studies investigating the development of auditory involuntary attention use a visual or auditory active categorization task (Gumenyuk, Korzyukov, Alho, Escera, & Näätänen, 2004; Wetzel, Widmann, Berti, & Schröger, 2006) or a visual passive task where participants watch a silent movie (Bonmassar, Widmann, & Wetzel, 2020). In both cases, the participants are instructed to focus on the targets or the movie and to ignore auditory distractors. However, it is unclear whether the results obtained in such tasks generalize to real-life activities. The distraction of attention during typical everyday activities such as playing a game or reading a book is barely investigated in this framework, particularly in children, and will be the focus of the present study.

Selective attention abilities develop throughout childhood and are not yet matured in adolescence (Hoyer et al., 2021; Slobodin, Cassuto, & Berger, 2018; Thillay et al., 2015; Wetzel et al., 2019). Recent behavioral auditory distraction studies report significant changes in distraction control between the age of around 4–8 years that continue to develop further (Hoyer et al., 2021; Wetzel et al., 2019). Neurophysiological and imaging studies demonstrate the development of the neural network underlying attentional control throughout childhood and early adolescence (Konrad et al., 2005; Mahajan & McArthur, 2015; Rueda, Rothbart, McCandliss, Saccomanno, & Posner, 2005). The attention network makes a transition from functional yet immature systems supporting attentional functions in children to the more definitive networks in adults (Konrad et al., 2005; Rueda et al., 2005).

Neuronal activity evoked by distractor sounds can be measured with high temporal precision using Electroencephalography (EEG). We are interested in the extent in which the early processing of task-irrelevant sounds (standard and novel sounds) is affected by the type of activity. We therefore measure the P1 component that reflects stimulus encoding processes (Liégeois-Chauvel, Musolino, Badier, Marquis, & Chauvel, 1994) and the P2 component that has been associated to early classification and attention processes (Crowley & Colrain, 2004). Both components are prominent in the first years of life and diminish until adulthood (Wunderlich, Cone-Wesson, & Shepherd, 2006). The P3a component, associated with orienting of attention toward distractor sounds, can be already observed in toddlers (Putkinen, 2012) and is reliably observed in children in the age range of 5−10 years (Bonmassar et al., 2020; Gumenyuk, Korzyukov, Alho, Escera, & Näätänen, 2004; Määttä et al., 2005; Wetzel, Widmann, & Schröger, 2011).

Measures of brain oscillations provide another important means of understanding the underlying neural mechanisms of involuntary capture of attention by the irrelevant distractor. Sustained attention relies on theta oscillations, which organizes visual thalamic signaling during sustained attention (Clayton, Yeung, & Cohen Kadosh, 2015; Xie, Mallin, & Richards, 2018; Yu et al., 2018). Oscillatory activity in the theta band has also been linked with cognitive control (Cavanagh & Frank, 2014; Cooper et al., 2019; Ryman et al., 2018), working memory load (Hsieh & Ranganath, 2014; Jensen & Tesche, 2002), and auditory distractor processing (Fuentemilla, Marco-Pallarés, Münte, & Grau, 2008; Ishii et al., 2009; Ponjavic-Conte, Dowdall, Hambrook, Luczak, & Tata, 2012) in adults. Theta power has been shown to increase in response to deviant and novel sounds in a sequence of auditory tones (auditory oddball) in healthy adults (Fuentemilla et al., 2008; Hsiao, Wu, Ho, & Lin, 2009). On the other hand, continuous auditory distraction noise attenuates the activity of theta band oscillations and a higher distraction causes a greater reduction of theta band power and phase synchrony (Ponjavic-Conte et al., 2012). These findings suggest that examination of theta band activity might help to understand the dynamics of neural oscillations during auditory distraction processing and involuntary attention orienting. However, the relation between involuntary attention orienting and theta oscillations has not yet been investigated in children.

This study focuses on the impact of task-irrelevant sounds on neuronal mechanisms that drive perception and attention during three naturalistic activities in children aged 7-8 years and adults. We chose to focus on 7-8-year-olds for this study because the brain morphology and networks underlying visual and auditory attention control undergo developmental changes throughout middle childhood and shift towards maturation thereafter. Thus, the 7-8-year-old age group offers a unique opportunity to measure attention-related brain activity in children and for comparison with the matured brain activity in adults. Moreover, previous studies had demonstrated modulations of attention by task-irrelevant sounds in 7-8-year-olds (Bonmassar et al., 2020; Wetzel, Widmann, & Scharf, 2021). Thus, there is scope to study the effect of task-irrelevant sound processing during typical everyday activities in this age group.

In three conditions, we measured the ERP components P1, P2, P3a, and concurrent oscillations in the theta band from EEG in response to task-irrelevant standard and novel oddball sounds while participants played a card game, read a book or watched a silent movie. We addressed two main questions: 1) Do different activities, that are typical for children, differently modulate the processing of task-irrelevant sounds and involuntary attention mechanisms? 2) Does the type of activity differently modulate sound processing and attention in children and adults? A memory card game requires maintaining and modifications of task-relevant processes in the working memory while a silent movie, which has been often used in previous developmental studies, can be watched more passively. We expect that when playing the card game, fewer resources are available for processing and paying attention to task-irrelevant sounds. This is expected to be reflected by reduced amplitudes of the ERP components P1, P2, P3a in the playing compared to the watching condition. Indeed, previous studies found that an increased task-related working memory load reduced the distraction by task-irrelevant novel sounds in adults (Lv et al., 2010; SanMiguel, Corral, & Escera, 2008). Due to the ongoing development of working memory load in middle childhood (Gathercole et al., 2004), it is expected that task-irrelevant sound processing is more impaired during playing the card game in children than in adults. Moreover, reading is a highly automatic process in adults, and it is expected that more resources are allocated to process task-irrelevant sounds than in the playing condition. How strong children are distracted by sounds during reading in the second year of school is an open question since reading is not yet an automatic process as it is in adults (Ehri, 1995). We are also interested in general age effects on novel sound processing and attention. Children tend to show larger ERP amplitudes in response to task-irrelevant distractor sounds across multiple task domains (Bonmassar et al., 2020; Gumenyuk et al., 2004; Määttä et al., 2005; Wetzel, 2015). Consequently, we expect larger amplitudes of the ERP component P3a, which has been discussed to reflect increased orienting of attention and evaluation processes in children compared to adults. Due to the lack of studies, the modulation of theta power by novel sounds in children is still an open question. Moreover, studies with adults report inconsistent findings on an increase or decrease of theta with distraction (Fuentemilla et al., 2008; Hsiao et al., 2009; Ponjavic-Conte et al., 2012).

## 2. Materials and methods

### 2.1. Participants

A total of 38 participants (20 children, 18 adults) participated in this study. Data of two children were excluded from analyses due to technical problems during recording and one child did not finish the experiment. The mean age of the remaining 17 children (female=10) was 7.76 years (range = 7 years 10 months - 8 years 7 months) and the mean age of adults (female = 10) was 22.89 years (range = 18 years – 30 years 8 months). Participants and parents in the case of children confirmed that they had a normal or corrected-to-normal vision, no medication with effects on the central nervous system, and no history of attention-related disorders. Children gave oral assents, and their parents and adult participants gave written informed consent. Participants were rewarded with vouchers for a local toy shop (children), money, or course credits (adults) for their participation. The experimental protocol was approved by the local ethics committee.

### 2.2. Auditory Stimuli

Standard sounds were triangle wave sounds with 500 Hz base frequency and the 120 novel sounds were common environmental sounds (e.g. person sneezing, pig grunting selected from a commercial CD, 1111 Geräusche, Döbeler Cooperations). Sounds had a duration of 200 ms. All sounds were tapered-cosine windowed (10 ms rise- and 10 ms fall-time). The intensity was root mean square matched and presented at 59 dB sound pressure level.

### 2.3. Tasks, instructions and task-related material

In the (1) playing condition, the participant and examiner played a memory card game. The game comprises a deck of cards that have a picture on only one side and two cards having the exact same picture (N=19 pairs). Cards are placed randomly face-down on the table in a matrix, and the players take turns flipping one card to reveal the picture and then flipping another one, which they think is the matching card. If the cards match, they are removed from the table, and the same player can continue. If they do not match, they are returned face-down at the original position, and the next player can begin. Before the start of a game, two to three moves were practiced. Participants were asked not to speak during the game. The examiner was instructed to interact naturally and consistently with all children. This means the examiner appreciatively smiles and nods when the child has found a pair. In the case of a child participant, the examiner would always let the child win to avoid frustration. If the sound sequence was finished before all pairs were detected, the game was ended and the player with the most pairs wins. This was announced before the start of the game. Furthermore, the experimenter and the participant agreed not to chat during the game and both stuck to it. The memory card game was identical for both age groups. For each condition, we asked participants to rate their interest in the task and how effortful the task was on a 3-point Likert scale and performed likelihood ratio analyses to evaluate if there were any differences between the two groups. There were no differences between the two groups in interest in (*X*^2^ (2) = 4.456, p = .108) and effort or difficulty of (*X*^2^ (2) = 1.421, p = .491) playing the memory game. In the (2) reading condition, participants were instructed to silently, and without moving lips, read a passage from an age-appropriate book. Children had the choice between a book about friends or dragons. Adults read a popular science book about psychiatry. All participants were informed that they subsequently would be asked to describe the content of the read text. In a previous training phase, children were asked to silently read one sentence and to tell the examiner with their own words the meaning of the sentence. As in the playing condition, there were no differences at the level of interest (*X*^2^ (2) = 1.588, p = .452) or difficulty (*X*^2^ (2) = 00, *p* = 1) in reading the books between the two groups. In the (3) watching condition, participants passively watched a silent animated movie that tells the story of a sheep and his family on an iPad (Model: MC497FD, IOS: 5.1.1) that was placed on the table in front of them. Participants rated how interesting they found the movie. Both group did not differ at the level of interest (*X*^2^ (2) = 4.144, p = .126) or effort (*X*^2^ (2) =.684, p = .710) in watching the movie. In all three tasks, participants were informed that sounds would be presented and that those sounds were not relevant to the task. Participants were asked to ignore the sound sequence. The lack of differences in interest and perceived difficulty in all conditions demonstrate that we can exclude the explanation that the EEG results are due to interest or difficulty in the task.

### 2.4. Procedure

The experiment was conducted in an electrically shielded and acoustically attenuated cabin. Participants were seated at a table. Seating positions were kept the same during the entire experiment. The examiner was sitting together with the child in the cabin in all three conditions to avoid different potential effects of the nearness of another person. Sounds were presented via two loudspeakers (Bose Companion 2 series III Multimedia speaker system). After the familiarization of children with the lab and the cabin, the procedure was explained and all questions of participants were answered. To assess their reading abilities, children were administered the SLS1-4 reading screening (Mayringer & Wimmer, 2003). The mean reading score (M=29.94, SD=8.8) is consistent with the standardization sample of second graders (M=25, SD=8.7). Then the electrodes were fixed.

The auditory oddball sequence was presented while participants performed the three different types of tasks (playing a memory card game, reading a book, watching a movie). The three conditions were presented in a balanced order across subjects. For each condition, two blocks with 600 trials each (480 standards and 120 novels) were presented consecutively. The interval between the onsets of two sounds was 1000 ms. Each block lasted 5 minutes and each condition 10 minutes. Novel sounds were pseudo-randomly presented with a probability of 20% embedded in a sequence of repeated standard sounds of 80%. Each novel sound was presented only once per condition, i.e., three times throughout the experiment, and at least two standards preceded a novel sound.

### 2.5. EEG data recording and preprocessing

EEG was recorded using BrainAmp MR amplifier by Brain Products GmbH (Gilching, Germany) at a digitization rate of 500 Hz and referenced to the tip of the nose. Ag/Ag-Cl Electrodes were placed at the following positions: FP1, FP2, FC1, FC2, FC5, FC6, F3, F4, F7, F8, T7, T8, C3, C4, CP1, CP2, CP5, CP6, P3, P4, P7, P8, O1, O2, Fz, Cz, Pz and at the left (M1) and right (M2) mastoids. Horizontal and vertical eye movements were measured by three electrodes that were placed at the left and right of the outer canthi, and below the left eye. EEG data preprocessing was performed with MATLAB software using the EEGLAB toolbox (Delorme & Makeig, 2004). Data were filtered offline with a 0.1 Hz high-pass filter (Hamming windowed sinc FIR filter, order = 8250, transition bandwidth = 0.2 Hz) and a 40 Hz low-pass filter (Hamming windowed sinc FIR filter, order = 166, transition bandwidth = 10 Hz) (Widmann, Schröger, & Maess, 2015). Data were segmented into 1.5 s epochs starting 500 ms before and ending 1000 ms after sound onset. Two standard trials at the beginning of each block and two standard trials after a novel trial were excluded from the analysis (Wetzel, 2015). Noisy channels were identified and removed from the data using the EEGLAB plug-in FASTER (Nolan, Whelan, & Reilly, 2010). To classify a channel as artifactual, FASTER calculates three parameters - variance, mean correlation and Hurst exponent - for each channel. A channel whose data had a Z-score of +/-3 for a parameter was deemed to be artifacted. These channels were removed from the analysis and interpolated after ICA and artifact rejection. This procedure removed on average 1.90 channels per subject (STD = 1.06, range = 0 to 5).

To further remove ocular artifacts and generic noise, independent component analysis (ICA) was performed on an identical copy of the dataset. To achieve an improved ICA decomposition, the copied dataset was high pass filtered at 1 Hz (Hamming windowed sinc FIR filter, order = 1650, transition bandwidth = 1 Hz) and epochs exceeding unusually large amplitude (+/-600 μV) were removed before ICA (Klug & Gramann, 2020). This procedure removes large non-stereotypical artifacts but keeps stereotypical artifacts such as blinks and eye movements to be later removed using ICA. After ICA decomposition, independent components (ICs) or ICA weights were then transferred from the ICA copied dataset to the original dataset. All further analyses were performed on this original dataset. Artifactual ICs were identified by a semiautomatic process that included using the IClabel plugin (Pion-Tonachini, Kreutz-Delgado, & Makeig, 2019) of EEGLAB and also visual inspections of individual ICs. IClabel identified ICs with eye artifacts such as eye blinks, saccades and muscle artifacts. Artifactual ICs as identified by IClabel and visual inspections were then removed from the original dataset. Each epoch was baseline corrected by subtracting the mean amplitude of the -200 to 0 ms window preceding sound onset. A voltage threshold rejection (+/-150 μV) was then applied to further exclude artifacts-laden epochs. After artifact rejection, missing channels were interpolated using spherical interpolation as implemented in EEGLAB. Table 1 presents the number of artifact-free trials retained after preprocessing for novel and standard sounds in each condition.

**Table 1.** The average number of trials for novel and standard sounds after preprocessing of EEG data in each condition for children and adult groups. Mean values are rounded to the nearest integers.

|  | Playing |  | Reading |  | Watching |  |
| --- | --- | --- | --- | --- | --- | --- |
|  | Novel | Standard | Novel | Standard | Novel | Standard |
| Children | 108 | 212 | 114 | 227 | 115 | 227 |
| Adult | 117 | 231 | 118 | 233 | 115 | 229 |

### 2.6. Principal component analysis (PCA)

In addition to computing conventional ERPs, we performed temporal principal component analysis (PCA) on our data. PCA is useful to identify the constituent components of the ERP and is particularly recommended for developmental ERP data due to the increased level of noise (Dien, 2010). In line with the current study, recent studies (Bonmassar et al., 2020; Dercksen, Widmann, Schroger, & Wetzel, 2020; Barry et al., 2020; Dien, Spencer, & Donchin, 2003) used temporal PCA on children’s and adult’s data to isolate ERP components associated with novelty response. PCA was performed using the MATLAB-based ERP PCA toolkit (Dien, 2010). For the PCA analysis, the length of the preprocessed epochs was reduced to 800 ms (-200 to 600) from the 1.5 s. PCA was computed using Geomin rotation (ε = 0.01) with a covariance relationship matrix and no weighting. The number of components to be retained was determined using Empirical Kaiser Criterion (Braeken & van Assen, 2017). PCA was computed on the individual averages of the standard trials and novel trials. We performed separate PCA for each group, and extracted 27 components from the ERPs of the adult group and 21 components from the ERPs of the children group. The PCA analysis was performed to obtain four components of interest namely P1, P2, and two subcomponents of the P3a, the early P3a (eP3a) and late P3a (lP3a; Escera et al., 1998; Yago et al., 2003). We analyzed the component peak amplitudes for all conditions and groups at the following electrodes: Fz (P1), Cz (P2), Cz (eP3a) and Fz (lP3a). The selection of electrode positions is based on existing literature (Bonmassar et al., 2020; Escera, Alho, Schröger, & Winkler, 2000; Winkler, Denham, & Nelken, 2009; Wunderlich et al., 2006). Table 2 presents the component numbers and variance explained by each component in adult and children groups.

**Table 2.** Component numbers and variance explained by each component in adult and children groups. The component time courses were rescaled to microvolts (μV) for each condition for statistical comparisons between the conditions and groups.

|  | Adults |  | Children |  |
| --- | --- | --- | --- | --- |
|  | Component | Variance (%) | Component | Variance (%) |
| P1 | 14 | 1.25 | 8 | 5.32 |
| P2 | 3 | 9.52 | 7 | 5.80 |
| eP3a | 1 | 20.89 | 3 | 13.38 |
| lP3a | 6 | 6.22 | 1 | 15.97 |

### 2.7. Time-frequency analysis

Time-frequency analysis was performed by the Matlab script provided by Cohen (2014). Time-frequency power was calculated for the epoched data. Time-frequency power provides a two-dimensional (time x frequency) estimate of average changes in spectral power (in dB) relative to the baseline. To compute time-frequency power, each epoch was convolved with complex Morlet wavelets, which estimated power in the frequency range 1–40 Hz (in 50 linearly spaced steps). The wavelet cycles were set at 2 cycles at the lowest frequency (1 Hz) increasing to 7 Hz at the highest frequency (40 Hz). Spectral power was calculated for all channels and separately for three conditions: playing, reading and watching, and also for both groups. In each condition, spectral power was estimated for novel sounds, standard sounds and novel minus standard difference. Power was calculated for each epoch relative to -200 to 0 ms pre-stimulus baseline period. Previous studies found that auditory oddball processing induced the strongest time-frequency effects at the Fz and Cz electrodes (Cahn, Delorme, & Polich, 2013). Therefore, the primary activity of interest was theta (4-7 Hz) rhythm oscillations at Cz and Fz electrodes.

### 2.8. Statistical Analyses

Statistical analysis was performed using the software JASP (Version 0.14.1; JASP Team, 2020). In order to compare the components between groups, the component time courses were rescaled to microvolts (μV) by multiplying the component loading with the standard deviation (SD) of component time scores (Dien, 2012). The resulting time course reflects the portion of the recorded waveform accounted for by each component scaled to μV, which is statistically comparable between children and adult groups. The statistical comparison was performed on the mean amplitude of +/− 20 ms time window around the peak of the component time course. The statistical analysis was based on the difference amplitudes computed by subtracting the mean standard-related ERP amplitude from the mean novel-related ERP amplitude (novel ERP minus standard ERP) of the components early P3a and late P3a observed in all conditions (Escera et al., 2000). We performed a Condition (playing, reading, watching) x Group (children, adults) ANOVA with Condition as the within-subject factor and Group as the between-subject factor for early P3a and late P3a components. We performed a separate Condition (playing, reading, watching) x Sound type (novel, sound) x Group (children, adults) ANOVA for P1 and P2 components. Greenhouse–Geisser corrections of degrees of freedom were applied when appropriate. Furthermore, given our apriori hypothesis, we performed post hoc comparisons with Bonferroni corrections to examine the nature of the sound processing and attentional orienting across the task conditions.

To reveal the time-frequency window with significant modulation of power, we combined the time-frequency data from the two groups, the three conditions and both novel and standard sounds, and performed point-wise analyses on the combined data. The point-wise analysis tested the significant modulation of power against a null hypothesis of no change in power (represented by zero). The point-wise analysis computed non-parametric permutation tests against zero for each time point. It performed 1000 permutations with false discovery rate (FDR) correction and p = .05. The point-wise test was performed at Fz and Cz electrodes. This procedure highlighted statistically significant time-frequency intervals around 100−300 ms time window at the theta (4-7 Hz) band (Figure 1). The 100−300 ms time window overlaps with the ERP components time course. To compare the oscillatory activity across the task conditions and groups, mean power at 100−300 ms time window in theta band was analyzed at Cz and Fz electrodes. We performed a Condition (playing, reading, watching) x Sound type (novel, standard) x Group (children, adult) ANOVA with Condition and Sound type as within-subject factors and Group as between-subject factor. We performed two separate ANOVAs for power at Cz and Fz electrodes. Likewise for ERP components, we performed post hoc comparisons with Bonferroni corrections to examine the theta power across task conditions.

**Figure 1.**
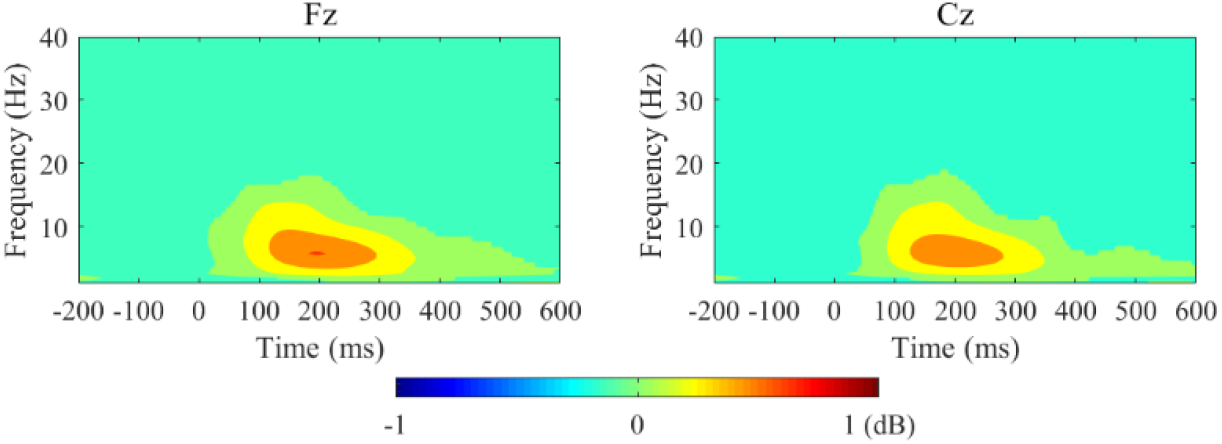
Time-frequency interval with masked non-significant activation in response to standard and novel sounds at Fz and Cz electrodes. A point-wise analysis was performed. The red area in the images indicates the statistically significant increase in time-frequency power. Green in images depicts non-significant (p > .05) time-frequency activity. The power (decibels, dB) of time-frequency activity is shown by the color bar. At both electrodes, there is a significant increase of power at the theta (4−7 Hz) band around 100−300 ms time window.

## 3. Results

### 3.1. P1

The peak latency of the P1 component was 90 ms in children and 78 ms in adults (Figure 2). The ANOVA showed a main effect of Group (*F*(1, 33) = 165.440, *p* < .001, *η*^2^ = .658) resulting from larger amplitude in children (*M* = 4.106) compared to the adults (*M* = .283). The analysis also revealed a significant main effect of Condition (*F*(2, 66) = 7.091, *p* = .002, *η*^2^ = .015) and a significant Condition x Group interaction (*F*(2, 66) = 7.102, *p* = .002, *η*^2^ = .015). There was no significant main effect of Sound type (*F*(1, 33) = .752, *p* = .392, *η*^2^ = .00008) or Sound type x Group interaction (*F*(1, 33) = 3.811, *p* = .059, *η*^2^ = .004). Post hoc comparisons for the Condition x Group interaction revealed that in the children group, reading (*M* = 4.376, *t* = 3.730, *p* = .006) and watching (*M* = 4.735, *t* = 5.068, *p* < .001) conditions had significantly greater amplitude compared to the playing condition (*M* = 3.373) whereas P1 amplitudes did not differ significantly in three conditions in the adult group.

**Figure 2.**
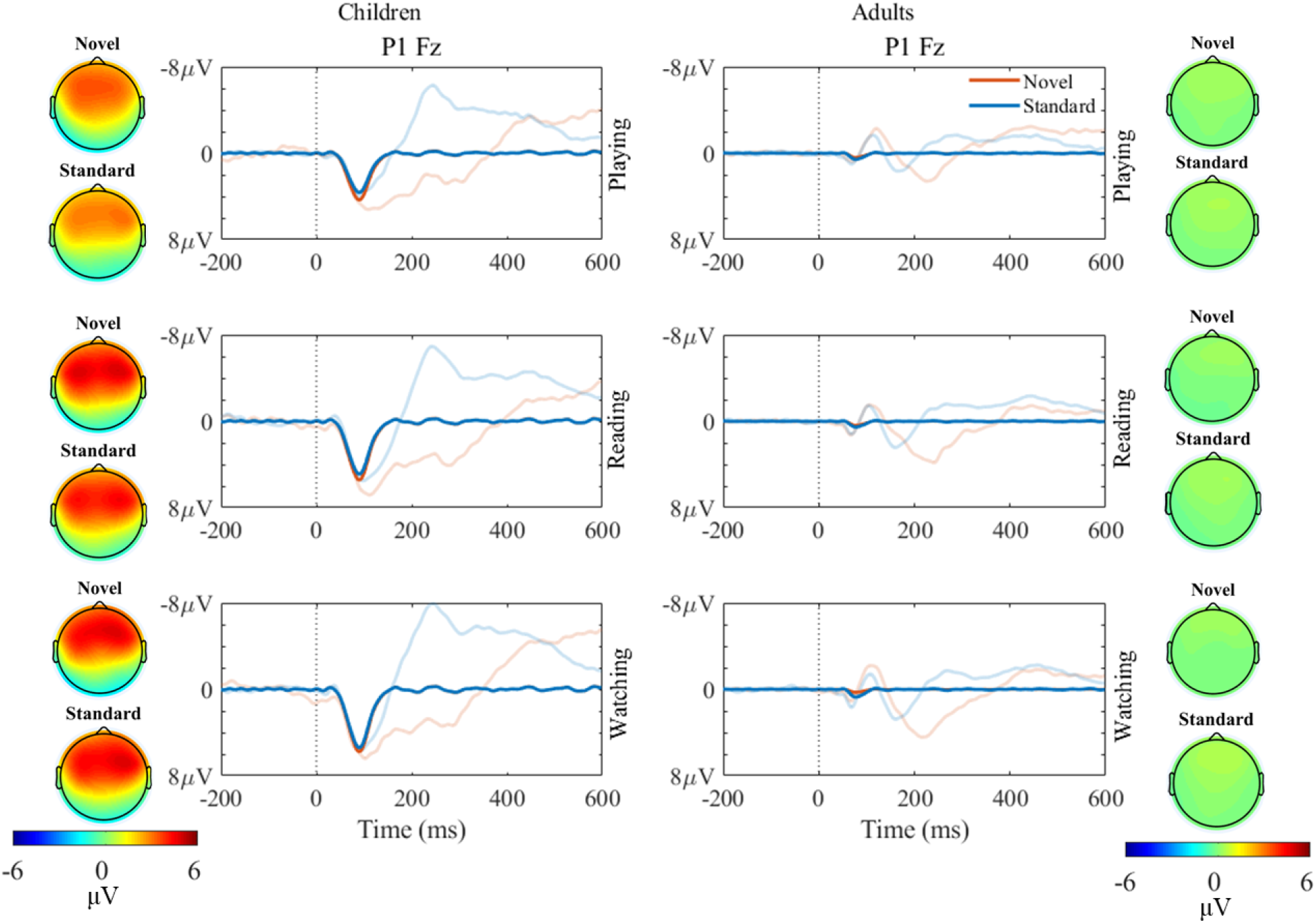
PCA components (opaque color) reflecting the ERP component P1 and topographies display the scalp distribution of the PCA components reflecting P1. The corresponding grand-averages ERPs at Fz are shown in transparent colors in the background. The left panels display PCA components and ERPs evoked by novel (red) and standard (blue) sounds in playing, reading and watching conditions in children. The right panels display PCA components and ERPs evoked by novel (red) and standard (blue) sounds in playing, reading and watching conditions in adults. Time 0 indicates the onset of the sounds. In children, the amplitudes of the P1 are smaller in the playing than in the watching and reading conditions. This difference was not observed in adults. No statistically significant difference was observed between amplitudes of novel and standard sounds.

### 3.2. P2

The peak latency of the P2 component was 158 ms in children and 166 ms in adults (Figure 3). The ANOVA did not reveal a significant main effect of Group (*F*(1, 33) = 1.901, *p* = .117, *η*^2^ = .024). The analysis showed a main effect of Condition (*F*(2, 66) = 5.524, *p* = .006, *η*^2^ = .038) but no significant Condition x Group interaction (*F*(2, 66) = 2.371, *p* = .101, *η*^2^ = .016). Post hoc comparisons for the main effect of Condition indicated that across sound types and groups reading (*M* = 1.987, *t* = 2.977, *p* = .012) and watching (*M* = 1.921, *t* = 2.768, *p* = .022) conditions evoked greater amplitude compared to the playing (*M* = 1.040) condition. The main effect of Sound type was not significant (*F*(1, 33) = .462, *p* = .713, *η*^2^ = .00004) but there was significant Sound type x Group interaction (*F*(1, 33) = 17.311, *p* < .001, *η*^2^ = .056). Post hoc comparisons for the Sound type x Group interaction showed that the two groups did not differ for standard sounds but children (*M* = 2.485) had significantly larger amplitude than the adults (*M* =.739, *t* = 3.100, *p* = .019) for the novel sounds.

**Figure 3.**
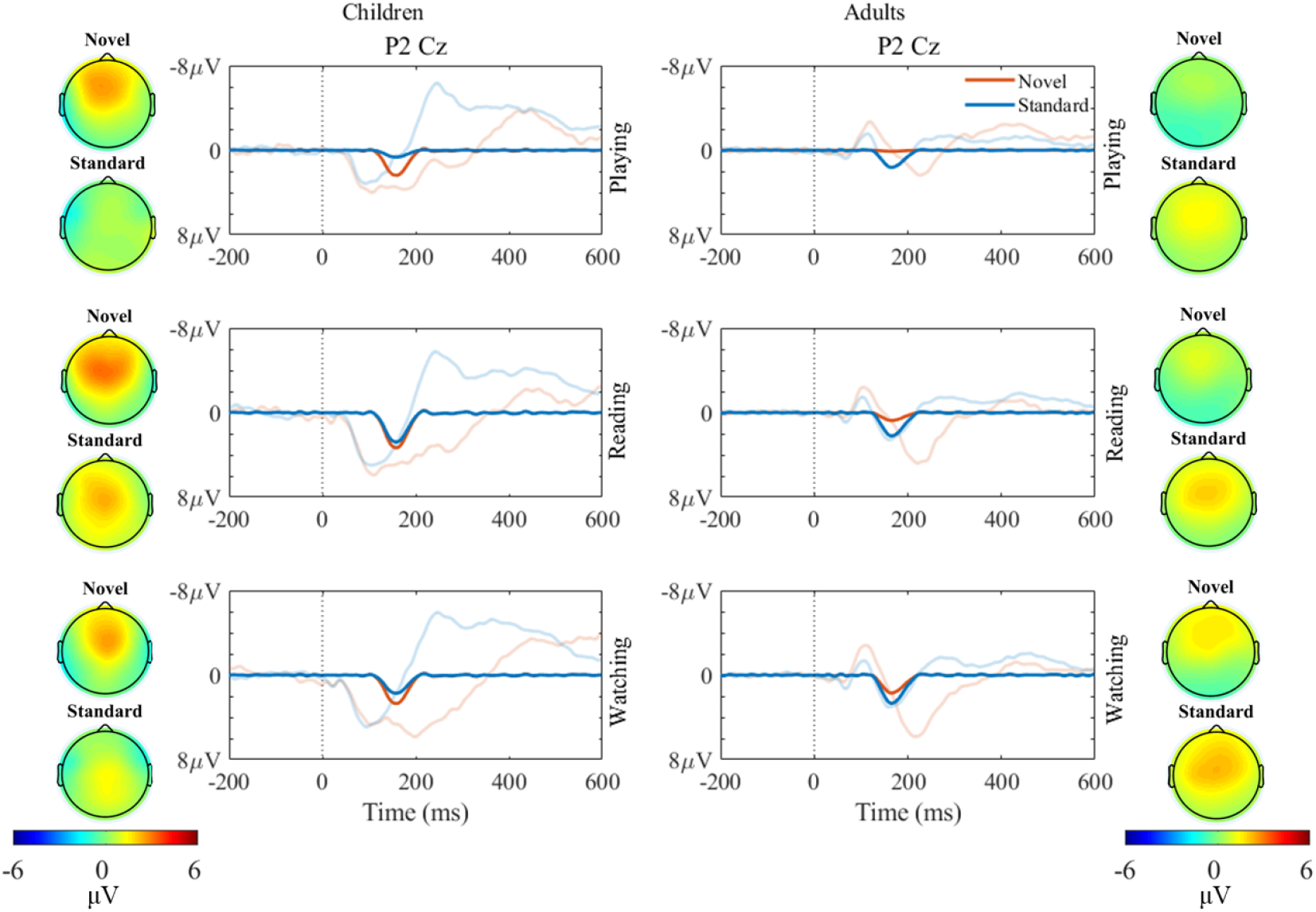
PCA components (opaque color) for the ERP components early P2 and topographies display the scalp distribution of the PCA components. The corresponding grand-averages ERPs at the Cz are shown in transparent colors in the background. The left panels display PCA components and ERPs evoked by novel (red) and standard (blue) sounds in playing, reading and watching conditions in children. The right panels display PCA components and ERPs evoked by novel (red) and standard (blue) sounds in playing, reading and watching conditions in adults. Time 0 indicates the onset of the sounds. In both age groups, P2 amplitudes were smaller in the playing than in the watching and reading conditions. P2 amplitudes were larger in response to the novel sounds in children than in adults, while they did not differ between groups in response to the standard sounds.

### 3.3. Early P3a (eP3a)

In the children group, the eP3a peak latency was 238 ms and the peak amplitude was descriptively maximum in the watching condition (*M* = 7.679) followed by the reading condition (*M* = 6.926) and minimum in the playing condition (*M* = 5.552). In the adult group, the eP3a peak latency was 238 ms and as in children, the peak amplitude was maximum in the watching condition (*M* = 5.341) followed by the reading condition (*M* = 4.924) and minimum in the playing condition (*M* = 2.849). The ANOVA revealed a significant main effect of Group (*F*(1, 33) = 11.024, *p* = .002, *η*^2^ = .143) resulting from larger amplitudes in the children (*M* = 6.676) compared to the adults (*M* = 4.338). The analysis also showed a significant main effect of Condition (*F*(2, 66) = 10.386, *p* < .001, *η*^2^ = .102). The Condition x Group interaction was not significant (*F*(2, 66) = 0.200, *p* = .819, *η*^2^ = .002). Post hoc comparisons showed that across groups watching (*M* = 6.477, p < .001) and reading (*M* = 5.892, *p* = .003) conditions resulted in significantly higher amplitude than the playing (*M* = 4.152) condition. The amplitude difference between reading and watching conditions was not significant (Figure 4).

**Figure 4.**
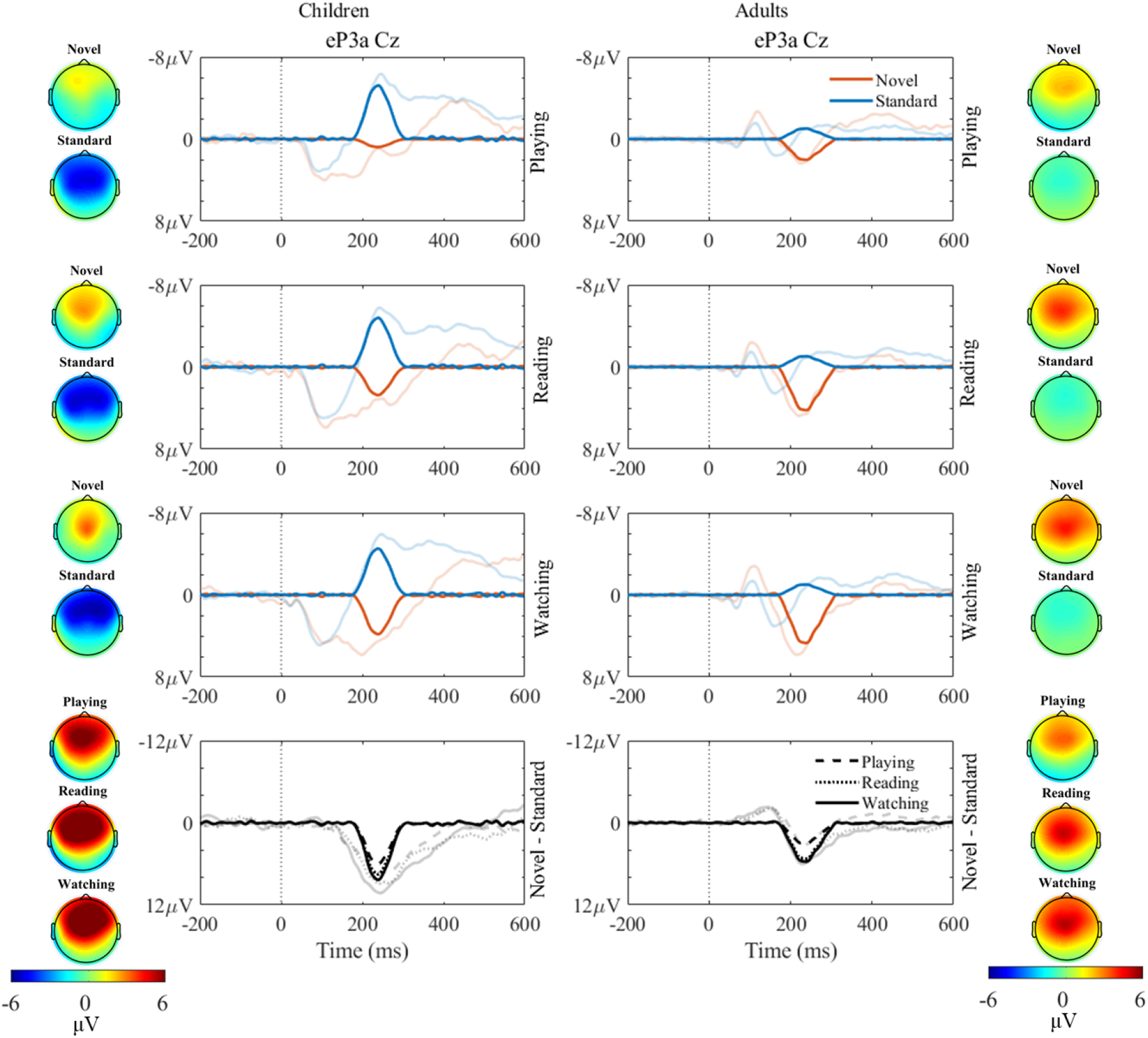
PCA components (opaque color) for the ERP components early eP3a and topographies display the scalp distribution of the PCA components. The corresponding grand-averages ERPs at Cz are shown in transparent colors in the background. The left panels display PCA components and ERPs evoked by standard sounds (blue), novel sounds (red), and novel minus standard for playing (dashed black), reading (dotted black) and watching (solid black) conditions in children. The right panels display PCA components and ERPs evoked by standard sounds, novel sounds, and novel minus standard for playing, reading and watching conditions in adults. Time 0 indicates the onset of the sounds. Early P3a amplitudes (novel – standard) were larger in children than adults.

### 3.4. Late P3a (lP3a)

The late P3a peak latency was 302 ms in children and the peak amplitude was descriptively maximum in the watching condition (*M* = 6.970) followed by the reading condition (*M* = 6.742) and minimum in the playing condition (*M* = 5.788). In the adult group, lP3a peak latency was 296 ms and the peak amplitude was maximum in the watching condition (*M* = 1.688) followed by the reading condition (*M* = 1.221) and minimum in the playing condition (*M* = 0.792). The ANOVA revealed a main effect of Group (*F*(1, 33) = 28.444, *p* < .001, *η*^2^ = .388) resulting from larger amplitudes in the children (*M* = 6.429) compared to the adults (*M* = 1.159). The analysis did not show a main effect of Condition (*F*(2, 66) = 2.301, *p* = .108, *η*^2^ = .010) or a Condition x Group interaction (*F*(2, 66) = .142, *p* = .868, *η*^2^ = .00006) (Figure 5).

**Figure 5.**
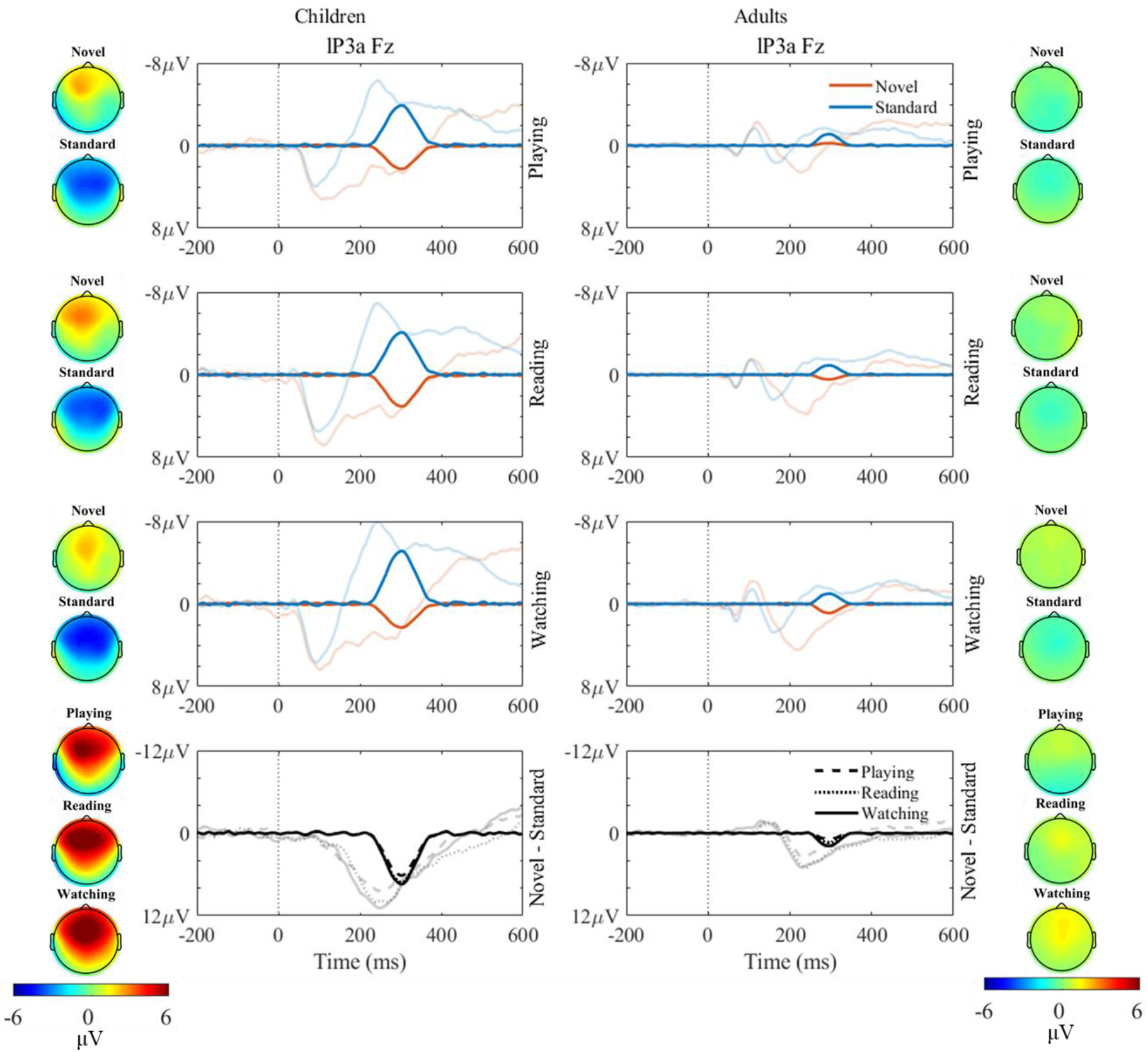
PCA components (opaque color) for the ERP components early lP3a and topographies display the scalp distribution of the PCA components. The corresponding grand-averages ERPs at Fz are shown in transparent colors in the background. The left panels display PCA components and ERPs evoked by standard sounds (blue), novel sounds (red), and novel minus standard for playing (dashed black), reading (dotted black) and watching (solid black) conditions in children. The right panels display PCA components and ERPs evoked by standard sounds, novel sounds, and novel minus standard for playing, reading and watching conditions in adults. Time 0 indicates the onset of the sounds. Late P3a amplitudes (novel – standard) were larger in children than adults.

### 3.5. Theta power at Cz electrode

Figure 6 shows theta power at the Cz electrode. For theta power at the Cz electrode, the ANOVA revealed a main effect of Group (*F*(1, 33) = 15.161, *p* < .001, *η*^2^ = .102) resulting from greater theta power in adults (*M* = .702) compared to the children (*M* = .257) across all conditions. There was a significant main effect of Condition (*F*(2, 66) = 6.208, *p* = .003, *η*^2^ = .031) and significant Condition x Group (*F*(2, 66) = 6.356, *p* = .003, *η*^2^ = .031) and Condition x Sound type (*F*(2, 66) = 3.801, *p* = .027, *η*^2^ = .017) interactions. The analysis also showed a significant main effect of Sound type (*F*(1, 33) = 55.648, *p* < .001, *η*^2^ = .129) and Sound type x Group interaction (*F*(1, 33) = 34.769, *p* < .001, *η*^2^ = .080). Post hoc comparisons for Condition x Group interaction showed that theta power did not differ between the three conditions in children (*M*^*P*^ = .273, *M*^*R*^ = .155, *M*^*W*^ = .322, all *p >* .05). However, in the adult group, reading (*M* = .798, *t* = 3.615, *p* = .009) and watching (*M* = .919, *t* = 4.635, *p* < .001) conditions had significantly greater power compared to the playing condition (*M* = .371). Post hoc comparisons for Sound type x Group interaction revealed that theta power between standard and novel sounds did not differ in children (*M*^*N*^ = .302, *M*^*S*^ = .198, *t* = 1.090, *p* = 1.000) but in adults novel sounds evoked highly significantly greater theta power than the standard sounds (*M*^*N*^ = 1.144, *M*^*S*^ = .247, *t* = 9.582, *p* < .001). Post hoc comparisons for Condition x Sound type showed that for the novel sounds, the watching condition had highly significantly greater power (*M* = .949, *t* = 4.266, *p* < .001) than the playing condition (*M* = .452). The reading condition also tended to had higher power than the playing condition (*M* = .788, *t* = 2.890, *p* = .068). The power did not differ in three conditions for the standard sounds. There was no significant Condition x Sound type x Group interaction (*F*(2, 66) = 1.299, *p* = .280, *η*^2^ = .006).

**Figure 6.**
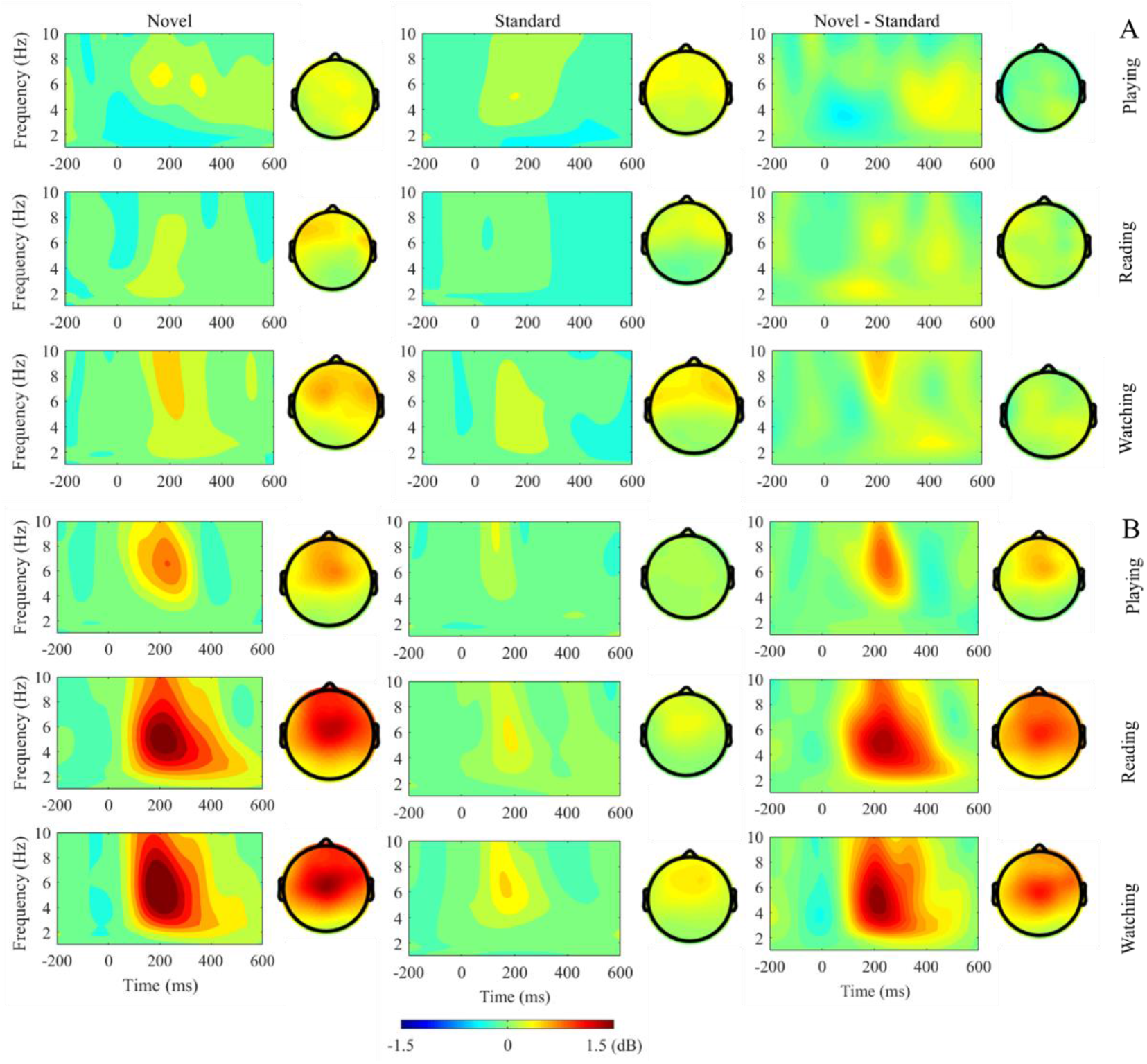
(A-B) Time-frequency activities at the Cz electrode and topographic maps of theta power (4-7 Hz) evoked by standard sounds, novel sounds and the difference between novel and standard. (A) The top three panels display power for the playing, reading and watching conditions of the children group. (B) The bottom three panels display power for the playing, reading and watching conditions of the adult group. Time 0 indicates the onset of standard and novel sounds. Power (dB) activity is shown by the color bar. Novel sounds evoked statistically significantly increased theta power compared to standard sounds in adults but not in children. Novel sounds in the reading and watching conditions evoked increased theta power compared to the playing conditions in adults whereas in children power did not differ in the three conditions.

### 3.6. Theta power at Fz electrode

The ANOVA for theta power at Fz showed a main effect of Group (*F*(1, 33) = 6.970, *p* = .013, *η*^2^ = .041), which resulted from greater theta power in adults (*M* = .627) across conditions than the children (*M* = .329). There was a significant main effect of Condition (*F*(2, 66) = 4.780, *p* = .012, *η*^2^ = .031). Post hoc comparisons indicated that across groups watching condition (*M* = .621) had significantly greater theta power than the playing condition (*M* = .373, *t* = 3.042, *p* = .010) but power between playing and reading (*M* = .536, *p* = .149) as well as between reading and watching conditions (*p* = .903) did not differ significantly. Condition x Group interaction was not significant (*F*(2, 66) = 1.342, *p* = .268, *η*^2^ = .009). The analysis also revealed a significant main effect of Sound type (*F*(1, 33) = 34.264, *p* < .001, *η*^2^ = .121) and a significant Sound type x Group interaction (*F*(1, 33) = 17.763, *p* < .001, *η*^2^ = .063). Post hoc comparisons for Sound type x Group interaction showed that theta power between standard and novel sounds did not differ in children (*M*^*N*^ = .446, *M*^*S*^ = .332, *p* > .05) but in adults novel sounds evoked highly significantly greater theta power than the standard sounds (*M*^*N*^ = .972, *M*^*S*^ = .276, *t* = 7.223, *p* < .001). There was no significant Condition x Sound type x Group interaction (*F*(2, 66) = .097, *p* = .904, *η*^2^ = .00005).

**Figure 7.**
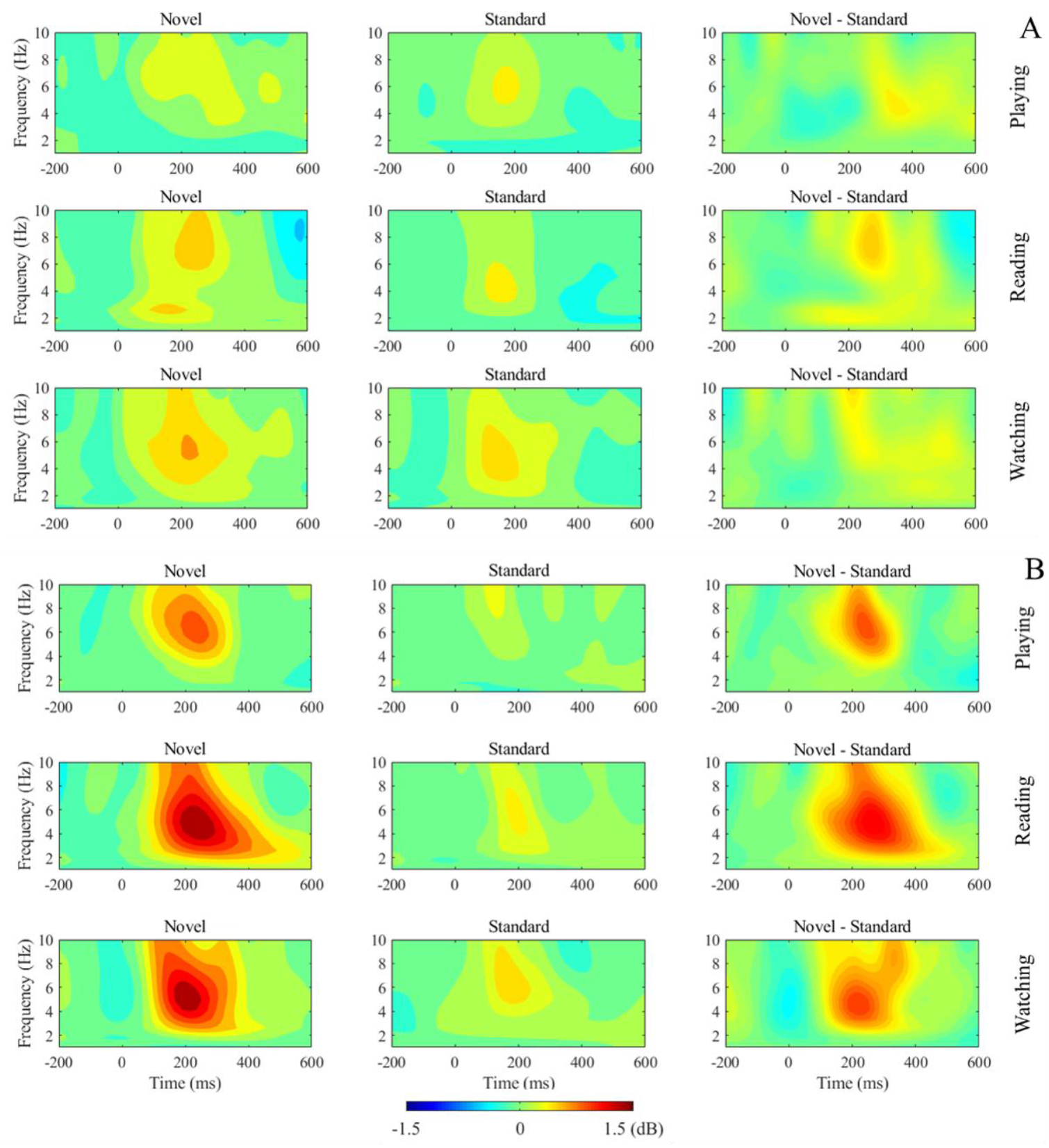
(A-B) Time-frequency activities at the Fz electrode at theta band (4-7 Hz) evoked by standard sounds, novel sounds and the difference between novel and standard. (A) The top three panels display power for the playing, reading and watching conditions of the children group. (B) The bottom three panels display power for the playing, reading and watching conditions of the adult group. Time 0 indicates the onset of standard and novel sounds. Power (dB) activity is shown by the color bar. Novel sounds evoked statistically significantly increased theta power compared to standard sounds in adults but not in children.

## 4. Discussion

The present study investigated the impact of different activities typical for children aged 7−8 years on the perception and attention of task-irrelevant sounds. We focused on three activities – playing a card game, reading a book, watching a movie, which children typically do at home and school in high-income countries. We investigated how these activities modulate brain responses to task-irrelevant, frequently repeated standard sounds and rarely and randomly presented novel sounds. Sound-related brain activity was investigated by the analyses of ERP components as well as oscillations from EEG data. Effects of the type of activity were observed on both ERP components and frontocentral theta power. The amplitudes of the ERP-components P2 and a subcomponent of the P3a, associated with stimulus processing and attention allocation, were reduced during playing compared to reading or watching in both age groups. In children only, the amplitude of the ERP component P1, reflecting early encoding processes, was reduced during playing. Sound-related theta power at the vertex was also reduced during playing in adults while children showed no effect of activity on theta power. Moreover, age modulated all ERP components of interest (P1, P2, eP3a, lP3a) as shown by increased amplitudes in children compared to adults. This age effect was specific for novel sounds for the P2 component. Fronto-central theta power in response to standard sounds was similar in children and adults but was considerably increased in response to novel sounds in adults compared to children.

### 4.1. The impact of activity on perception and attention of task-irrelevant sounds

Effects of activity on sound processing were observed already within the first 100 ms after sound onset. The amplitude of the P1 component was larger in the watching and reading condition than in the card playing condition in children. P1 reflects the early sensory processing of auditory stimuli presumably generated in the auditory cortex (Herrmann & Knight, 2001; Liégeois-Chauvel et al., 1994). Larger P1 amplitudes are specifically associated with enhanced processing of auditory stimuli (Rao, Zhang, & Miller, 2010; Tang et al., 2013). Results demonstrate increased neuronal activity in response to auditory stimuli in children when they read a book or watched a movie compared to when they played a memory card game with a human partner. Results can be interpreted as enhanced processing of task-irrelevant auditory information during reading and watching compared to playing in the children group. Such an impact of the activity on early encoding was not observed in adults indicating developmental differences in the susceptibility of encoding task-irrelevant stimuli during performed activities. Since no effect of sound type (standard vs novel) was observed it can be concluded that the encoding of all sounds, independent of their novelty, was modulated by the type of activity in children.

The temporally following positive ERP component, the P2 was also larger in the reading and writing condition than in the playing condition. The P2 is thought to be generated by multiple sources mainly located in the temporal lobe and partly driven by the output of the mesencephalic reticular activating system (Crowley & Colrain, 2004; Näätänen & Picton, 1987; Ponton, Eggermont, Kwong, & Don, 2000). The function of the underlying mechanisms is not yet well understood. It has been discussed that the P2 reflects stimulus classification processes (García-Larrea, Lukaszewicz, & Mauguiére, 1992). Following the current interpretation of the P2, increased amplitudes in the reading and watching condition compared to the playing condition indicate enhanced classification processes of task-irrelevant sounds in those conditions. Our results suggest that this influence of activity was similar in children and adults. Similar to earlier encoding processes, the type of activity modulated the processing of all sounds, independent of their novelty.

The type of activity affected also involuntary attention processes reflected by the P3a component. In line with previous information processing steps, playing the card game reduced P3a amplitudes compared to reading a book or watching a movie. This was observed for both age groups but only for the early part of the P3a. Both parts of the P3a, the centrally peaking early part and the frontally peaking late part are discussed to be involved in the attentional processing of distractor sounds (Escera, Alho, Winkler, & Näätänen, 1998; Yago, Escera, Alho, Giard, & Serra-Grabulosa, 2003). However, both components might not reflect exactly the same processes as they can be differently modulated by experimental conditions (Barry & Rushby, 2006; Escera et al., 2000; Horvath, Winkler, & Bendixen, 2008; Masson & Bidet-Caulet, 2019; Polich, 2007) and are assumed to be generated by partly different cerebral sources (Escera et al., 2000). However, the specific functional roles of the two P3a components are not yet completely understood.

ERP results indicate that playing a memory card game results in less processing and attending of task-irrelevant auditory information than reading a book or watching a movie. This impact of activity was similar in children and adults with the exception, that early sound encoding mechanisms are affected in children but not in adults. The differential impact of type of activity on neuronal mechanisms driving perception and attention might be explained with a different amount of claimed cognitive resources when performing the task. The processing of task-irrelevant sounds requires cognitive resources. It has been shown in previous auditory oddball studies that the presentation of task-irrelevant and unexpected novel sounds frequently impairs performance of the primary tasks in children (Wetzel et al., 2019) and adults (Escera et al., 2000). Assuming that the information processing capacity (Welford, 1952) and the attentional resources (Pashler, 1994) are limited, increased processing of task-irrelevant sounds could indicate reduced allocation of cognitive resources to the task at hand (playing, reading, watching). Or the other way around, reduced processing of the irrelevant sounds might indicate enhanced demand of cognitive resources by the primary task. Indeed, the playing condition might require enhanced working memory load compared to reading and watching conditions. In the playing condition, participants had to memorize the position and the covered pictures of the cards and have to update this information after each move. In addition, this was the only condition where participants interacted with a human partner. This might additionally require attentional resources for social perception and social evaluation processes (Capozzi & Ristic, 2018). The hypothesis of effects of increased task-related cognitive load on task-irrelevant sound processing is supported by EEG studies reporting reduced amplitudes of sound-related ERP components associated with early information processing and attention ((N1, P2, P3) during a difficult vs. easy gaming task in adults (Miller, Rietschel, McDonald, & Hatfield, 2011). This was interpreted by the authors as reduced attention allocation to the task-irrelevant sound sequence under high load conditions. This is consistent with studies reporting reduced amplitudes of the P3a component, that is discussed to reflect orienting of attention, when the working memory load was high (Harmony et al., 2000; Lv et al., 2010; SanMiguel et al., 2008). These previous studies demonstrate that ERPs evoked by task-irrelevant sounds can be a reliable marker of the workload in a task at hand (Allison & Polich, 2008). However, we did not directly measure the allocation of attention or the working memory load to the task. To avoid frustration during the memory card game, the experimenter let the child always win. Thus, no objective measurement of working memory performance was possible. This limitation of the present study could be addressed in future studies that directly control task performance and working memory load, for example by using n-back tasks.

We examined the theta power in three different conditions in children and adults. The results showed that the theta power did not differ in the three conditions in children. However, in adults, theta power was significantly reduced in the playing condition compared to the reading and watching conditions. Theta power reduction plays an important role in the maintenance of information in working memory (Brzezicka et al., 2019; Meltzer et al., 2007). Frontal theta power reduction was found to be associated with the increasing working memory load. In cognitive tasks, an increased working memory load reduced the theta power in pre- and lateral-frontal areas (Brzezicka et al., 2019; Meltzer et al., 2007). Individuals who were faster in accessing working memory showed a stronger decrease in theta power (Brzezicka et al., 2019). Our findings of reduced theta power in the playing condition are in line with these previous studies. An increased working memory load has been proposed to control attention and thereby shield against distraction in the auditory modality (Berti & Schröger, 2003; SanMiguel et al., 2008). The reduction of central theta power especially in novel trials in the playing condition thus indicates a reduced attentional allocation to task-irrelevant sounds because of an increased working memory load.

The reduction of ERP amplitudes and theta power in the playing condition might be a marker of greater working memory load during task performance. To our knowledge, the effect of working memory load on brain oscillations during auditory distractor processing has not yet been investigated. Given the lack of previous findings, our results should be interpreted with caution since we did not systematically manipulate the working memory load in this study. Nonetheless, our results do illustrate different modulations of brain responses as a function of processing task-irrelevant sounds during typical everyday activities in children and adults. The present results offer insights for future research investigating the relationship between brain activity and auditory involuntary attention capture as a function of working memory load during different activities.

### 4.2. Immature neuronal networks underlying novel processing

In line with our hypothesis, we consistently observed effects of age on task-irrelevant sound processing in the ERP components P2 and P3a and in frontal-central theta power. While the P2 amplitude in children in response to standard sounds did not differ from those of adults, the P2 amplitudes evoked by novel sounds were larger in children than in adults. This indicates that the mechanisms underlying P2 such as categorization processes (García-Larrea et al., 1992) are sensitive to the novelty of task-irrelevant sounds in children. This sensitivity is in line with previous studies reporting attention-related influences on the P2 in the context of motivationally significant stimuli, for example, target stimuli or new stimuli, that can evoke increased amplitudes of the P2 (Bonmassar et al., 2020; Getzmann, Wascher, & Schneider, 2018; Potts, 2004). The increased sensitivity of children to novel sounds, observed in this study, is in line with behavioral data that show that children in this age range are particularly prone to the novelty of distracting sounds (Wetzel, Widmann, & Scharf, 2021). The temporally subsequent P3a component was also affected by age. Novel sounds evoked a different activity in the time range of the P3a component (early and late P3a) compared to standard sounds in both age groups. That is, the task-irrelevant sound sequence is further processed on a level that enables that novel sounds capture involuntarily attention. This difference was larger in children than in adults. This has been shown previously and was interpreted as increased orienting of attention toward novel sounds in children (Bonmassar et al., 2020; Gumenyuk et al., 2004).

While there is a number of ERP studies (Kushnerenko, Van Den Bergh, & Winkler, 2013; Näätänen, Petersen, Torppa, Lonka, & Vuust, 2017; Wetzel, 2014), there is limited research with primary school age children investigating oscillatory activity during an auditory oddball paradigm. Oscillatory activity in the theta frequency band has been associated with modulation of attention (Cahn et al., 2013; Harris, Dux, Jones, & Mattingley, 2017; Missonnier et al., 2006; Yordanova, Rosso, & Kolev, 2003). Theta power in frontal and central regions increases substantially in response to auditory novel or target stimuli compared to standard or non-target stimuli in adults (Hsiao et al., 2009; Ishii et al., 2009; Missonnier et al., 2006). In this study, the children group showed a significantly reduced theta power than the adults for novel sounds. While adults showed significantly increased theta power in response to novel sounds than to standard sounds, children did not show an increase in the theta response for the novel sounds. This was observed in all conditions. Caravaglios et al. (2010) examined theta power in an auditory oddball paradigm in patients with mild Alzheimer’s disease (AD) and healthy elderly controls. They observed no significant post-stimulus theta power increase for either target or non-target auditory tones in the AD group whereas in healthy controls an enhancement of post-stimulus theta power was found only during target tone processing. Caravaglios et al. (2010) argued that the lack of theta enhancement during target tone processing indicated an impairment of attentional resource allocation and a deficit of the attentional/working memory system. This pattern of lack of theta power increase for auditory target stimuli has been also observed in adults with schizophrenia (Doege et al., 2009). Doege et al. (2009) suggested that the absence of theta power increase in subjects with schizophrenia is a manifestation of impaired stimulus evaluation, memory retrieval, and a lack of sustained attention. Previous studies have reported that children possess an immature attentional system and the selective attention abilities do not mature until adolescence (Bonmassar et al., 2020; Hoyer et al., 2021; Thillay et al., 2015). In this study, the lack of theta power enhancement for the novel sounds in children may be the result of deficits in stimulus evaluation and sustained attention due to immature neuronal systems of auditory attention. This is in line with the attention networks (alerting, orienting, executive) described by Petersen and Posner (2012) and their developmental trajectories. Both the orienting as well as the executive network significantly develop throughout middle childhood and interact with each other (Pozuelos, Paz-Alonso, Castillo, Fuentes, & Rueda, 2014). These immature attention systems contribute to an increased level of distractibility, which might be the reason for the absence of theta power increase in children. This age-related difference in theta power was observed only for novel sounds but not for standard sounds indicating that the underlying mechanisms are specific for novel sound processing and sensitive to maturational changes.

## 5. Conclusion

The current study extends the knowledge on the processing of task-irrelevant auditory information and attentional allocation in childhood. The oscillation analyses showed that the theta band power might reflect neuronal processes underlying involuntary attention orienting caused by task-irrelevant auditory stimuli. The special focus of this study was on the effects of everyday activities on the perception and attention of task-irrelevant sounds. Our findings demonstrate that the type of activity affects the processing of surrounding task-irrelevant information on both, the pre-attentive level as well as on higher attention levels. Playing a memory card game, which is hypothesized to require increased working memory load, reduced the processing and attention of task-irrelevant sounds. These effects differ between children and adults on the level of stimulus encoding and in terms of oscillatory activity on the level of attention. Our results might have implications for developmental research investigating the influence of cognitive demands during different activities and attention allocation to a variety of auditory distractors.

## Acknowledgements

We are grateful to all children and families who participated in our study and to Andreas Widmann for support in programming the experiment.

## Funding

The study was supported by the German Research Foundation (WE/5026-1-2), the Center for Behavioral Brain Sciences Magdeburg financed by the European Regional Development Fund (ZS/2016/04/78120) and the Leibniz Association (P58/2017).

## Data availability statement

The data are available from the authors on request through emailing the corresponding author. The presented manuscript uses open source Matlab toolbox and software for EEG and statistical analyses, which are cited throughout the manuscript.

## Conflict of Interest

The authors declare no conflict of interest or known competing financial interests or personal relationships that could have appeared to influence the findings reported in this paper.

## Notes

### Competing Interest Statement

The authors have declared no competing interest.

### Summary of Updates

The manuscript has been revised after comments and suggestions from the reviewers. The updates include a more detailed description of the task and an additional figure for the time-frequency power at the Fz electrode. The figure legends have been edited to make them clearer. Furthermore, the discussion section of the manuscript has been also expanded.

